# *In silico* exploration of potent flavonoids for dengue therapeutics

**DOI:** 10.1101/2024.03.24.586491

**Authors:** Anuraj Phunyal, Achyut Adhikari, Jhashanath Adhikari Subin

**Affiliations:** Central Department of Chemistry, Tribhuvan University, Kirtipur, Kathmandu, Nepal; Bioinformatics and Cheminformatics Division, Scientific Research and Training Nepal Private Limited, Kausalttar, Bhaktapur, Nepal

**Author notes:** These authors contributed equally to the work (Anuraj Phunyal, Jhashanath Adhikari Subin).

**Keywords:** ADMET, binding free energy, DENV-2, molecular docking, molecular dynamics, RMSD, RMSF

## Abstract

Dengue poses a persistent and widespread threat with no effective antiviral drug available till now. Several inhibitors have been developed by targeting the viral non-structural proteins including methyl transferase (NS5) of the dengue virus with possible therapeutic values. In this work, virtual screening, molecular docking, molecular dynamics simulations (200 ns), and assessments of free energy changes to identify potential candidates from a database of flavonoids (*ca.* 2000) that may have good curative potential from the disease. The binding affinity of flavonoids, namely Genistein-7-glucoside (FLD1), 6’–O-Acetylgenistin (FLD2), 5,6-dihydroxy-2-(4-hydroxyphenyl)-7-[3,4,5-trihydroxy-6-(hydroxymethyl)oxane-2-yl]oxychromen-4-one (FLD3), Glucoliquiritigenin (FLD4), and Chrysin-7-O-glucoronide (FLD5) showed the binding affinities of −10.2, −10.2, −10.1, −10.1, −9.9 kcal/mol, respectively, and possessed better values than that of the native ligand with showed (−7.6 kcal/mol) and diclofenac sodium (−7.3 kcal/mol). Drug-likeness of these compounds was acceptable and no toxicity was hinted by ADMET predictions. The stability of the protein-ligand complexes was accessed from 200 ns molecular dynamics simulation in terms of various geometrical parameters; RMSD, RMSF of residues, Rg, SASA, and H-bond of the protein-ligand complexes. The binding free energy changes of these compounds were calculated by the MM-PBSA solvation model with negative values (less than −38.01±7.53 kcal/mol) indicating the spontaneity of the forward reaction and favorability of the product formation. The geometrical and thermodynamic parameters infer that the flavonoid binds at the orthosteric site of the target protein of DENV-2 and could inhibit its functioning resulting in the prevention of the disease. Overall, this study highlights the anti-DENV activity of five flavonoids, positioning them as promising candidates for further development as antiviral agents against dengue infection.

## 1. INTRODUCTION

The Dengue virus (DENV) belongs to the Flaviviridae family [1] and is estimated to impact approximately 400 million individuals worldwide annually according to epidemiological evidence [2]. The economic impact causes dengue *ca.* US$8.9 billion annually [3]. Dengue virus causes dengue fever, dengue hemorrhagic fever (DHF), and dengue shock syndrome (DSS) [4]. Dengue fever exhibits signs in individuals, including a rise in body temperature reaching 40°C, discomfort in muscles and joints, intense headaches, reddening of the face, the appearance of skin rashes, and severe flu-like indications [5]. During hemorrhagic fever, a rapid increase in body temperature is observed, when entering the critical phase, along with a notable decline in both albumin and cholesterol levels [6]. The escape of plasma during fever in individuals infected with dengue is a critical condition that can result in dengue shock syndrome [7]. Currently, there is a lack of specific antiviral medications for treating dengue infection. Sanofi-Pasteur’s tetravalent dengue vaccine (CYD-TDV, Dengvaxia) has been approved for clinical use in several countries [8,9]. It has raised concerns regarding its overall efficacy in the general population [10,11]. Therefore proper, effective, and safe therapeutics for the treatment and cure of dengue are still lacking.

DENV has four serotypes (DENV-1, DENV-2, DENV-3, DENV-4), among them DENV-2 is also called severe dengue and is commonly in Asia and Latin America [12–15]. The single-stranded positive-sense RNA genome is its genetic material, approximately 11 kb in size. The genomic RNA consists of three main sections: a 5’ untranslated region, a single open reading frame (ORF), and a 3’ untranslated region. ORF contains the genetic code for a polyprotein, which undergoes processing by both viral and host protease [16]. Genomic RNA translated to polyprotein produced three structural proteins capsid (C), pre-membrane (prM), and envelope (E) which are responsible for forming the different parts of the virion. Additionally, it encodes seven non-structural proteins (NS1, NS2A/B, NS3, NS4A/B, NS5) that play a role in viral RNA replication [17]. This article focuses on the NS5 protein of DENV-2 is a good therapeutic target [18]. NS5 protein is the largest, consisting of approximately 900 amino acid residues. It possesses two distinct domains: a methyl transferase domain (MTD) located at the N-terminal and an RNA-dependent RNA polymerase domain (RdRp) at the C-terminal (Fig 1).

**Fig 1.**
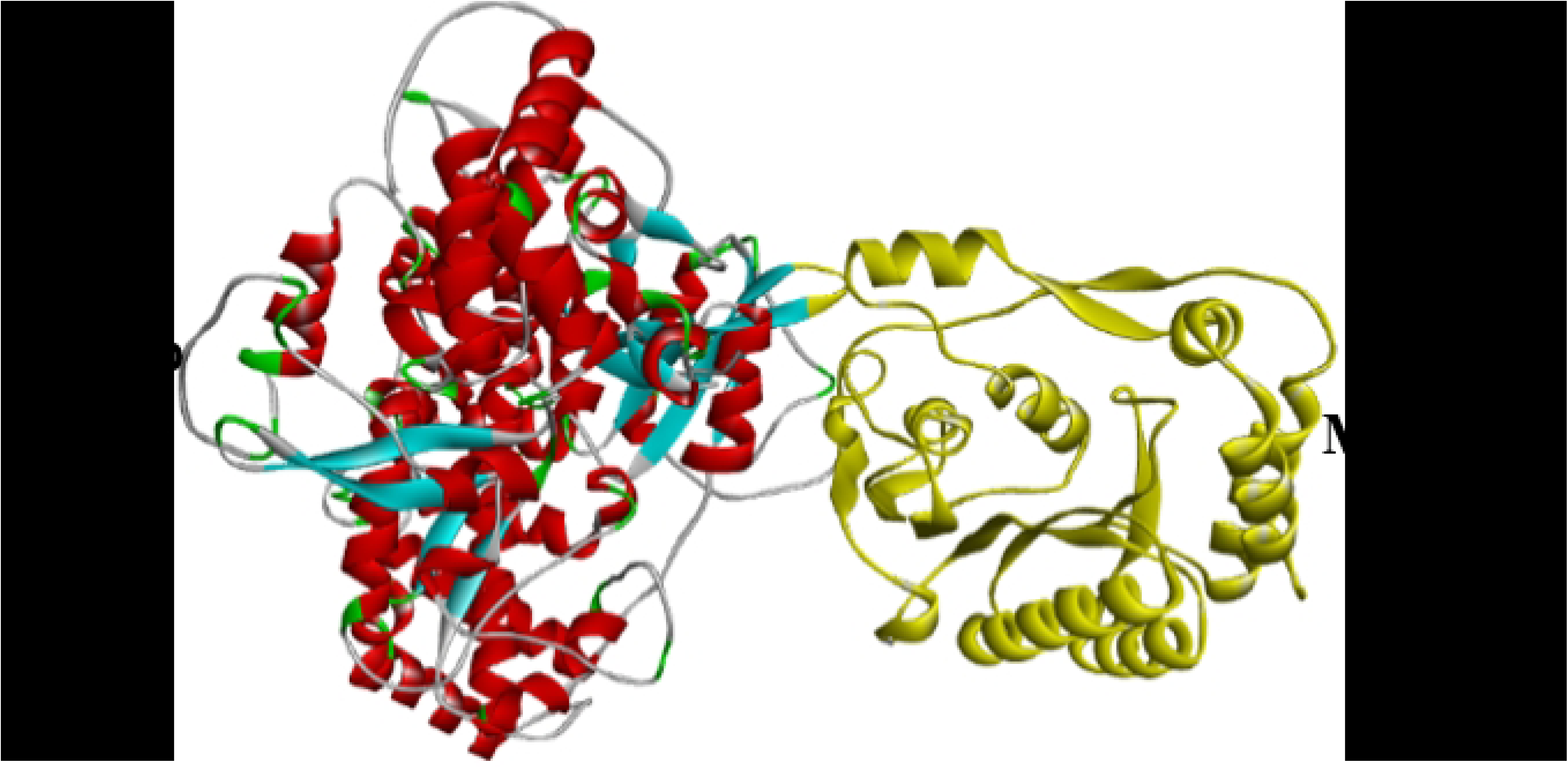
NS5 Protein of dengue virus.

The MTD of NS5 plays a crucial role in preventing viral mRNA degradation by 5’-exoribonucleases. It accomplishes this by adding a methyl group to the mRNA, which protects it from being recognized and degraded by cellular enzymes and makes sure that the eukaryotic translation initiation factor can recognize the mRNA [19]. At the GTP (guanosine triphosphate) binding site, the methylation occurs at the guanosine N-7 position, resulting in the formation of N-7-methylguanosine. Here, methylation takes place at the ribose, also leading to the formation of 2’ -O- methyl-adenosine. S-adenosylmethionine (SAM) is utilized as the source of methyl groups for this process [20–22]. It is a secondary metabolite present in virtually all bodily tissues and fluids, and plays a pivotal role in numerous vital functions, such as the immune system. MTD is a good target, competitive inhibition of SAM, and a good drug design option [23].

Medicinal plants have been found to contain a wide range of phytochemicals, including organosulfur compounds, limonoids, furyl compounds, alkaloids, polylines, coumarins, thiophenes, peptides, terpenoids, polyphenolics, and saponins. Flavonoids, which are a class of polyphenolic compounds, occur widely in various plants. They exist in different forms such as free compounds, glycosides, and methylated derivatives. Flavonoids have been reported to possess antiviral activity [24]. It has been chosen for this study to explore its ability to inhibit the function of NS5 methyl transferase of dengue virus and to cure the disease [25].

The use of natural substances derived from *Carica papaya* and *Euphorbia hirta* to treat dengue has been reported in recent studies [26]. Despite numerous *in vitro* studies conducted over the years, effective dengue inhibitors have yet to be discovered [27]. Computer-aided techniques are becoming more crucial in drug discovery because they help uncover potential therapeutic options quickly from a large pool of possibilities with low failure rates and recall [28]. Computational tools in drug discovery are an efficient and cost-effective way to determine and predict the therapeutic impact of various substances in a relatively shorter period than experimental analysis high throughput screening (HTS) [29,30]. To identify the potential therapeutics against the dengue virus, this work includes toxicity screening, drug-likeness prediction, molecular docking, ADMET prediction, molecular dynamics simulation (MDS), and binding free energy calculations. This work aims to identify NS5 inhibitors from flavonoids in low-cost environments with the least or null toxicity and optimum drug-likeness using computational assessment. The results would help in identifying the hit candidates that could be used for further *in vitro* and *in vivo* trials synergistically.

## 2. Computational experiment

### 2.1. Selection and preparation of ligand database

A dataset of ligands with anti-dengue properties [31] was constructed using a similar structure search and retrieved from the PubChem server [32] in SDF format. The ligands were then prepared for molecular docking by adding polar hydrogen using the Avogadro software (version 1.1) [33] after that energy minimized of the ligand structure using several parameters such as the conjugate gradient approach, energy convergence of 10^-8^ eV, and UFF force field for 2000 cycles were chosen for structural optimization. The minimization was performed through multiple attempts until no significant structural changes were observed (ΔE=0). The SDF format of ligands changed to pdb format by using Pymol (version 2.5.4) [34] program and further converted to pdbqt format followed by the addition of gasteiger charges. The bond order and the molecular formula were checked for the correct stoichiometry of the structure in the given compound.

### 2.2. Protein preparation and target structure validation

The NS5 protein of DENV, with the PDB ID of 5ZQK (resolution of 2.30Å, X-ray diffraction, expression organism: *E. coli*) was chosen as the target [35]. The crystal structure of this protein was obtained as a PDB file from the Protein Data Bank server (https://www.rcsb.org/). The chain A of NS5 protein was processed by using the Pymol program. First, water, followed by the addition of polar hydrogens then ions and other non-standard amino acid residues were removed. Furthermore, AutoDockTools (version 1.5.7) [36] was utilized to change the pdb form to pdbqt form of protein and incorporate Kollman’s charge to make the system neutral. For protein structure validation, the SAVES v6.0 web server (https://saves.mbi.ucla.edu/) with three modules namely, ERRAT [37], VERIFY3D [38], and PROCHECK [39] (Ramachandran plot) were used to identify protein sequences and assessed the quality of protein structure.

### 2.3. Homology modeling of receptor

Since the target protein was missing some amino acid residues, homology modeling was performed by using the SWISS-MODEL server [40]. For modeling of the target, the FASTA sequence was downloaded and the BLAST [41,42] database search method was employed, A template was generated, was evaluated using the Global Model Quality Estimation (GMQE) and Quaternary Structure Quality Estimation (QSQE) [43] For the selected template, a 3D protein model was generated. The generated model was subsequently utilized for molecular docking. The native ligand docking was carried out based on the orthosteric pocket reported in the crystal structure. Also, the CASTp server [44] was used to verify the location of the orthosteric site.

### 2.4. ADMET prediction

The database of *ca.* 2000 molecules was screened based on toxicity criteria from the ADMETlab 2.0 server [45], ProTox-`II`, and swissADME servers, for pharmacokinetics and pharmacodynamics studies [46,47]. To study ADMET properties of FDA-approved drugs (diclofenac sodium) repurposed for dengue [48], other hit candidates were comparatively analyzed and described.

### 2.5. Molecular docking

To find the potential candidates for dengue therapeutics molecular docking was performed using Autodock Vina software (version 1.1) was used for molecular docking [49]. The accessory program AutoGrid was used for generating the grid maps of protein. A standard grid box size of 40 Å × 40 Å × 40 Å, the grid center set at x, y, and z dimensions of (57.568, 3.995, and 47.337), the energy range of 4, spacing of 0.375 Å, and the number of modes of 20 with an exhaustiveness value of 32 were used. The structures were visualized using the Avogadro (version 1.1) program and verified in terms of geometry, stoichiometry, and bond order. The protein-ligand complex with the best binding affinity was saved in pdb format and used for molecular dynamics simulation and molecular-level analysis. Visualization of the binding interaction between protein and ligand was done through Biovia Discovery Studio 2021 Client software [50].

### 2.6. Molecular dynamics simulation (MDS)

The ligand-protein adduct was simulated by using the GROMACS program [51], employing the charmm27 force fields [52] for both the ligand and receptor. The ligand force field was obtained from the Swissparam server as .zip format [53]. To solvate the system, the TIP3P water model was used in a triclinic box, with a spacing of 12 Å at the sides chosen to minimize the spurious interactions between the periodic images [54]. The system was neutralized with counter ions and an isotonic solution of NaCl (0.15 M) was added. The equilibration was processed in four steps, each of 200 ps, at a physiological temperature of 310 K. The first two steps achieved NVT equilibrium, while the last two steps achieved NPT equilibrium. During equilibration, temperature coupling (modified Berendsen thermostat) [55], and pressure coupling (Berendsen isotropic) were applied, with PME (particle mesh Ewald) used for long-range Coulomb interactions [56]. In the final phase of the simulation, a production run was conducted for a duration of 200 ns without imposing any restraints with a step size of 2 fs. Geometrical parameters such as RMSD (Root Mean Square Deviation), RMSF (Root Mean Square Fluctuation), R_g_ (Radius of Gyration), SASA (Solvent Accessible Surface Area), and hydrogen bond count were calculated from the MDS trajectory using the inbuilt modules of the GROMACS program.

### 2.7. Binding free energy calculation by MM/ PBSA method

Molecular mechanics with the Poisson-Boltzmann surface area solvation (MM/PBSA) method was used in calculating the changes in binding free energies of the protein-ligand complex. The change in binding free energy of the protein-ligand complex is given by a linear equation [57]

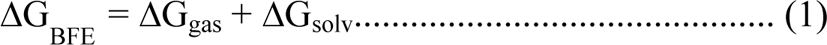

ΔG_gas_ is the sum of electrostatic (ΔE_EL_), and van der Waals energies (ΔE_VDWAALS_), and the solvation-free energy (ΔG_solv_) is the sum of polar (ΔE_PB_), and nonpolar (ΔE_NPOLAR_) parts [58]

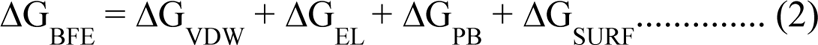

Based on the assessment of free energy changes, the forward reactions’s spontaneity and viability were assessed, 200 frames about the equilibrated part of the MDS trajectory were used in the calculation of the BFE.

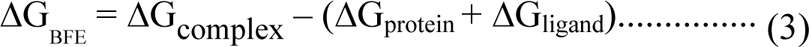

## 3. RESULTS AND DISCUSSION

### 3.1. Target 3D structure validation

Before molecular docking, the NS5 protein was prepared and validated for its structural integrity and robustness from server-based calculations. The overall quality factor from the ERRAT module of the protein was found to be 94.2197%, and from the VERIFY module, the structure of the protein was verified (87.02%). From the PROCHECK of the protein, a Ramachandran plot was obtained and showed over 90% of residue lies in the most favored region (Fig S1). The active site of protein contains SER56, GLY86, TRP87, LYS105, ASP131, VAL132, and ASP146, as key amino acid residues which were deposited in the protein data bank [59].

### 3.2. Docking protocol validation

Docking protocol validation was done through the docking of the native ligand at the active site resulting in an RMSD of less than 3 Å relative to the pose in the crystal structure (Fig 2). This justified the parameters taken and the algorithm used in obtaining the binding affinities and the ligand poses.

**Fig 2.**
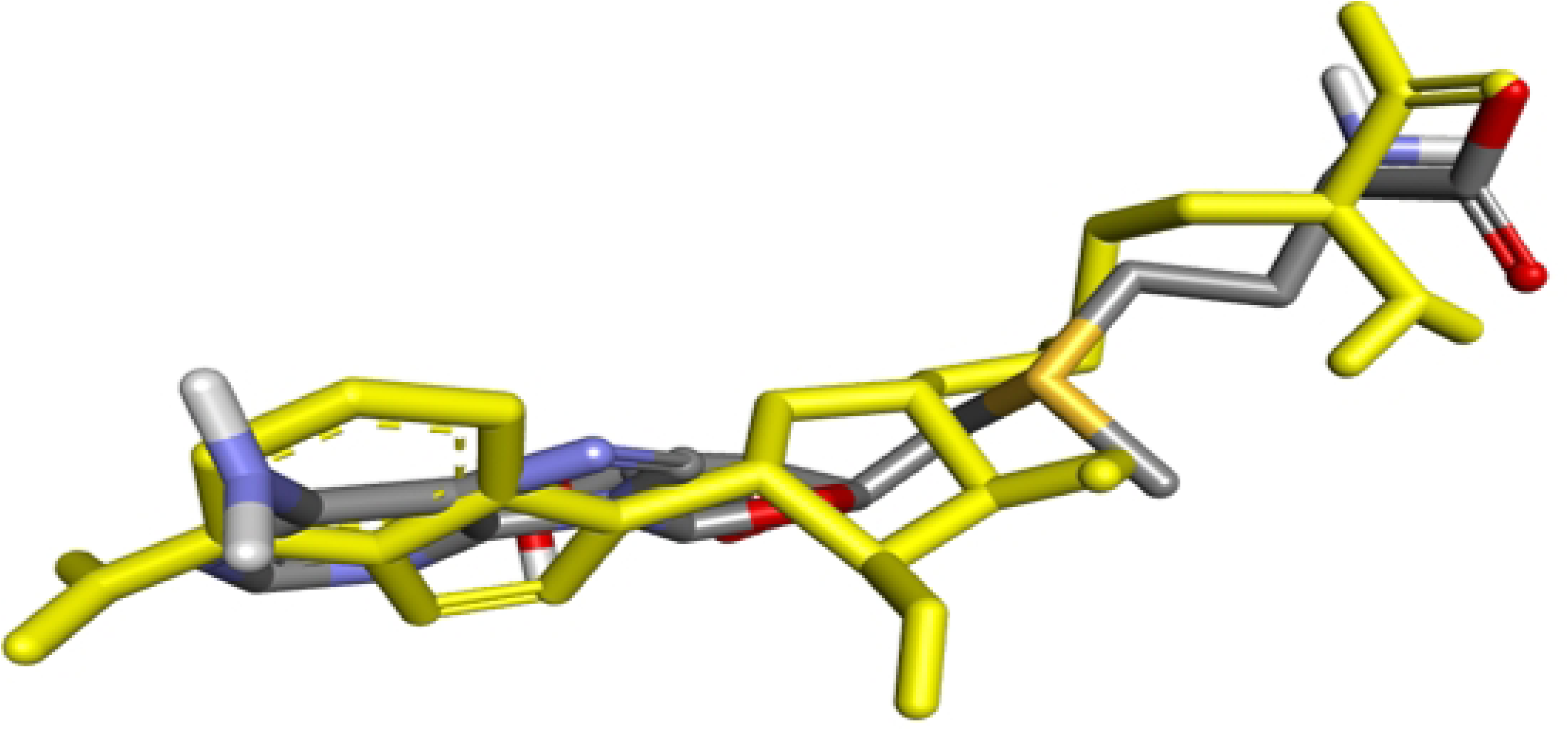
Superimposition of co-crystallized ligand (yellow) and docked ligand (Grey-red) with RMSD = 2.618 Å.

### 3.3. Analysis of modeling result of protein

A modified protein structure (PDB ID: 5ZQK) was created by modeling based on the template structure 5ZQK. Specifically, Model_01 was generated. The alteration in the amino acid residue numbering occurred due to the insertion of a missing amino acid in the template position [60]. The protein’s GMQE and QSQE values were determined to be 0.84 and 0.82 respectively, exceeding the threshold of 0.7. After modeling, a high-quality model of the template was constructed, with the active site residues identified as SER79, GLY109, TRP110, LYS128, ASP154, and VAL155. All the calculations were performed using the model generated by homology modeling.

### 3.4. ADMET analysis

Conducting an ADMET study can help reduce the likelihood of failure of drug candidates [77]. As a result, a comprehensive analysis was conducted subsequently to evaluate both drug-likeness and ADMET characteristics of the potential candidate. The hit candidates (class 5, LD50 >2500 mg/kg) in this study successfully passed assessments for hepatotoxicity, carcinogenicity, mutagenicity, immunotoxicity, and cytotoxicity (Table S1). It was found that the candidates exhibit plasma protein binding (PBB) of less than 90%, indicating a suitable PBB and high therapeutic index (Table S2). Furthermore, it demonstrated acceptable results (ranging from 0 to 0.3) for skin sensitization and minimal eye corrosion/irritation, almost reaching non-sensitizing levels (Table S3). The CNS permeability of the hit candidates was determined to be (logPS<-3), indicating impermeability to the central nervous system (Table S4). Additionally, the gastrointestinal (GI) effects of the hit candidates were found to be very low.

The hit candidates would not inhibit CYP2D6, CYP1A2, CYP2C19, CYP2C9, and CYP3A4, which play a vital role in metabolizing xenobiotics substances and minimizing the negative effects of drug interaction [78]. The hit candidates do not act as renal Oraganic Cation Transporter 2 (OCT2) substrate which is unlikely to have a significant impact on renal clearance. An FDA-approved drug, Diclofenac sodium (class 3, LD50 53 mg/kg) demonstrated a remarkably elevated plasma protein binding (PPB) rate of 99.21% and thus possessed a relatively low therapeutic index recognized for its potential to induce respiratory toxicity and provoke skin sensitization. The high gastrointestinal absorption exhibited by this drug enables effective system distribution and therapeutic effects but it also possesses hepatotoxic effects. Hence, the drug candidates showed lower toxicity compared to both reference ligand and reference drug. Especially, FLD1, FLD2, FLD3, FLD4, and FLD5 exhibited promising potential as effective candidates for dengue through inhibition of ene of its proteins. Overall studies showed the hit candidates exhibit favorable pharmacokinetic and pharmacodynamic properties.

### 3.5. **Molecular docking analysis**

In this study, investigated Thirty-four molecules that passed the ADMET screening were for further molecular docking. These selected molecules along with native ligands, and reference drugs, were evaluated against NS5 methyl transferase (chain A), where competitive inhibition was observed between proteins and flavonoids, consistent with recent research [61]. Among the 34 flavonoids studied, binding affinities ranging from −10.2 to −7.6 kcal/mol, surpassing those of the reference ligand (−7.6 kcal/mol) and reference drug (−7.3 kcal/mol) (Table 1), indicating higher binding efficiency and stability of the complexes [62]. FLD1, FLD2, FLD3, FLD4, and FLD5 emerged as the top 5 candidates (Fig 3) based on binding affinities.

**Fig 3.**
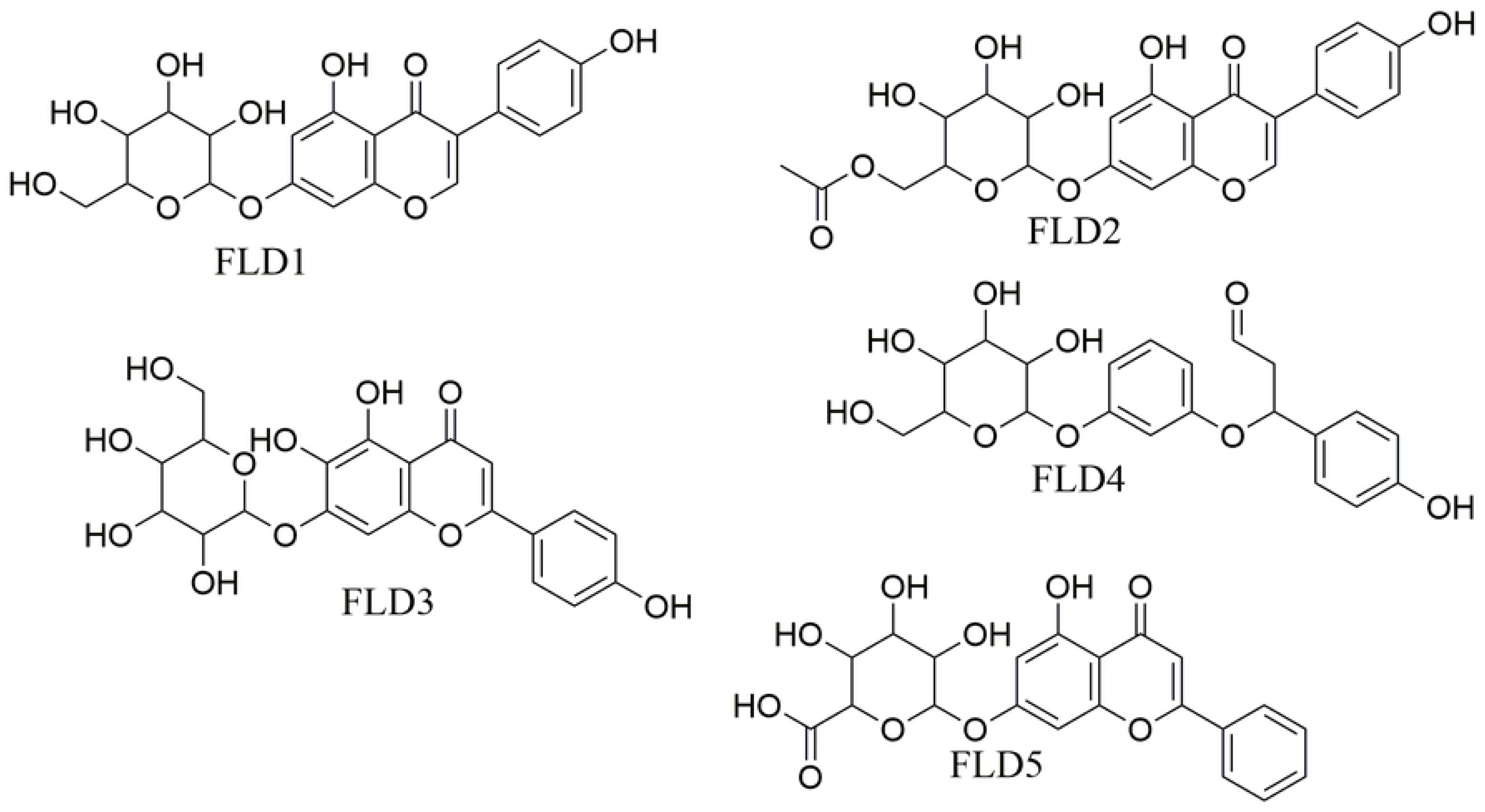
2D chemical structures of top five ligands (based on binding affinity)

**Table 1.**
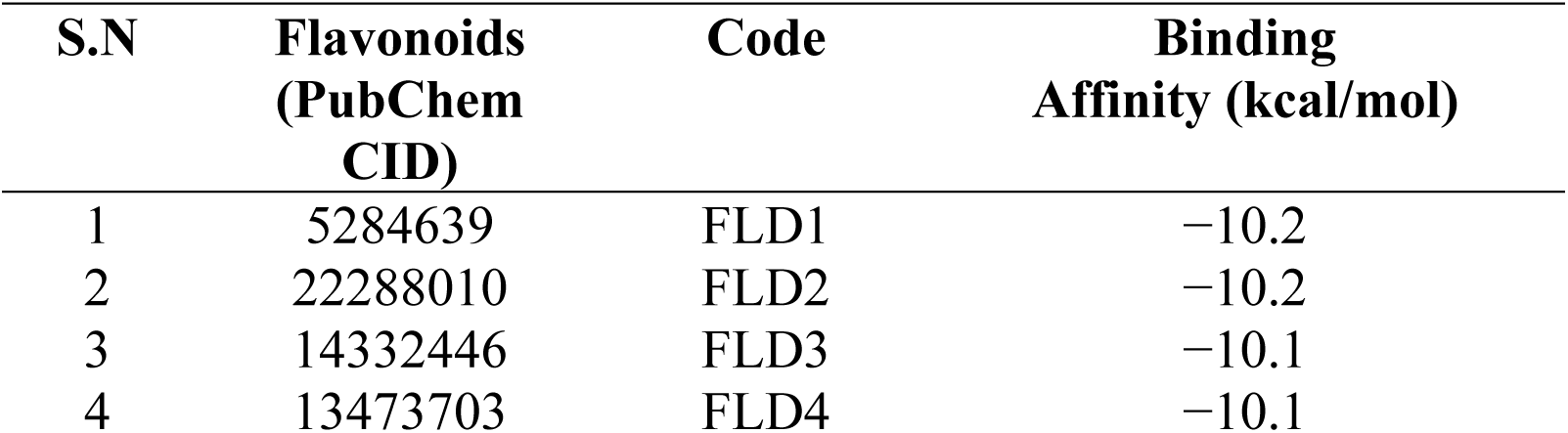
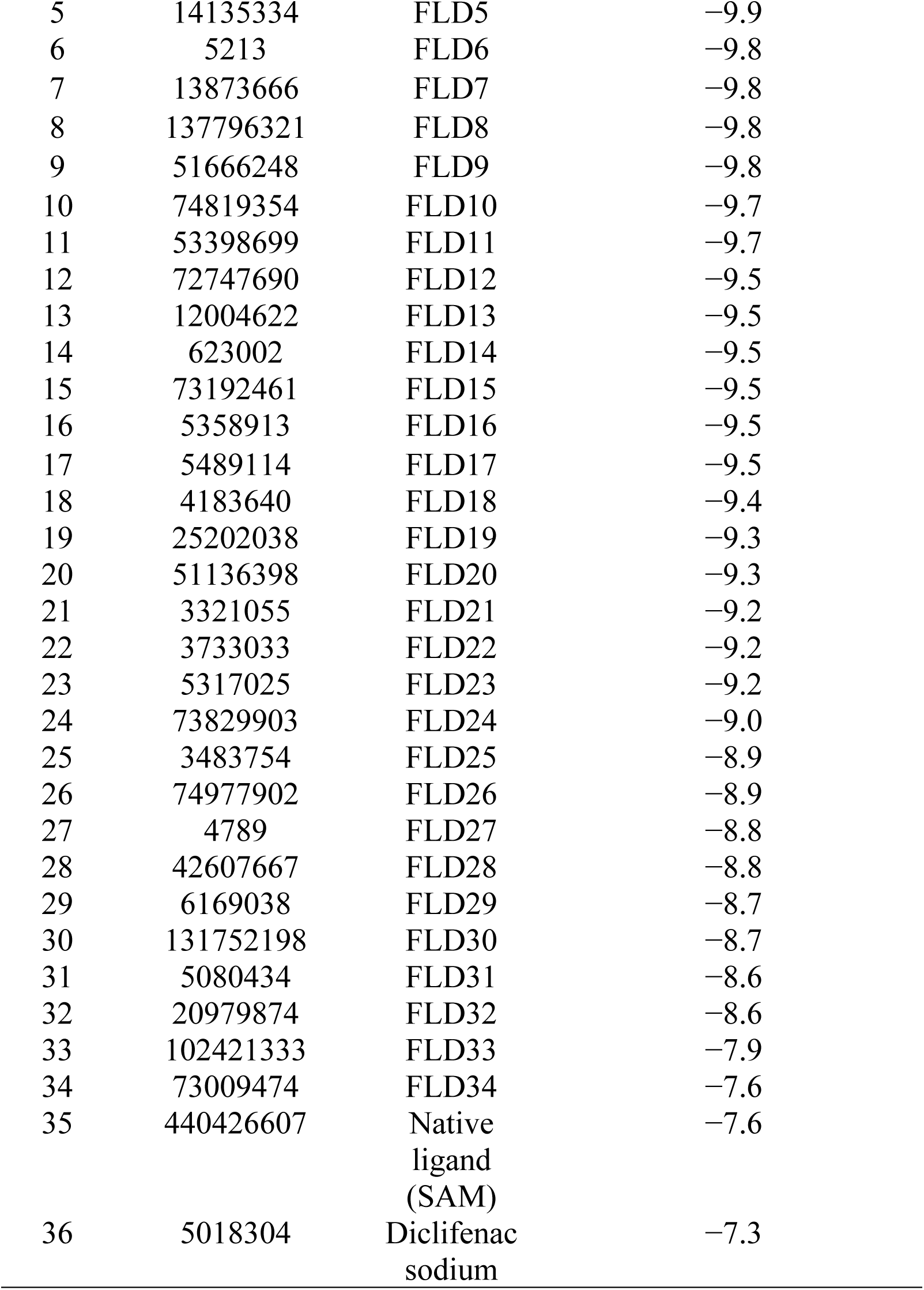
List of flavonoids with Binding Affinities from molecular docking calculations.

Notably, FLD1 and FLD2 show exemplary binding affinities of −10.2 kcal/mol each, and ranked highest among the molecules tested. FLD1 formed hydrogen bond interactions with GLY108 (2.35 Å), **GLY109** (2.19 Å), THR127 (2.37 Å), **LYS128** (2.94 Å), GLU134 (2.71 Å), and **ASP154** (2.13 Å), along with GLY81 (3.22 Å), HIS133 (3.80 Å), ARG107 (1.86 Å), **VAL155** (4.97 Å), and ILE170 (3.80 Å) are additional binding site residues and 9 van der Waals interactions are present. Similarly, FLD2 engaged in hydrogen bonding with GLY81 (2.82 Å), **GLY109** (2.85 Å), **TRP110** (2.39 Å), THR127 (2.29 Å), **LYS128** (2.90 Å), GLU134 (2.34 Å), other non-covalent interactions with GLY81 (3.71 Å), ARG107 (1.55 Å), **LYS128** (4.94 Å), **VAL155** (1.18 Å), HIS133 (3.03 Å), and ILE170 (3.63 Å) and 11 van der Waals interaction with NS5 in this complex. FLD3, FLD4 exhibited strong binding affinities of −10.1 kcal/mol with NS5 protein. FLD3, for instance, formed hydrogen bonds with **SER79** (2.97 Å), ARG107 (2.56 Å), GLY108 (2.28 Å), and GLY171 (2.18 Å), additional interactions were observed with **LYS128** (4.93 Å), **VAL155** (5.03 Å), and ILE170 (3.61 Å) amino acid residues and 16 van der Waals. Similarly, FLD4 and FLD5 engaged in bonding interactions with specific residues, and other non-covalent interactions, contributing to their binding affinities. The active site residues were engaged with all the protein-ligand complexes, which is the same as in previous studies with (same receptor) the distinction being a variation in residue number [63,64]. Despite the overall favorable interactions, some unfavorable interactions were present in FLD1, FLD2, FLD4, and FLD5 due to steric hindrance, incompatible geometry, electrostatic repulsion, and solvent effects [65]. Importantly, FLD1 (IC_50_ 100 nM) [66], FLD2, FLD3 (IC_50_ 10.6 µM) [67], FLD4, and FLD5 interacted through a combinations of hydrogen and hydrophobic interactions Fig 4(b), Fig 5(b), Fig 6(b), and Fig 7(b), and Fig 8(b) respectively. Analysis of interactions revealed distinct hydrophobic and hydrophilic regions Fig 4(a), Fig 5(a), Fig 6(a), Fig 7(a), and Fig 8(a), with the brown-colored regions indicating hydrophobicity and blue-colored regions indicating hydrophilicity. Furthermore, FLD1, FLD2, and FLD5 exhibited greater bonding interactions at the orthosteric site residues of the protein compared to FLD3 and FLD4 (Table 2), contributing to their dynamic stability after molecular docking [68].

**Fig 4.**
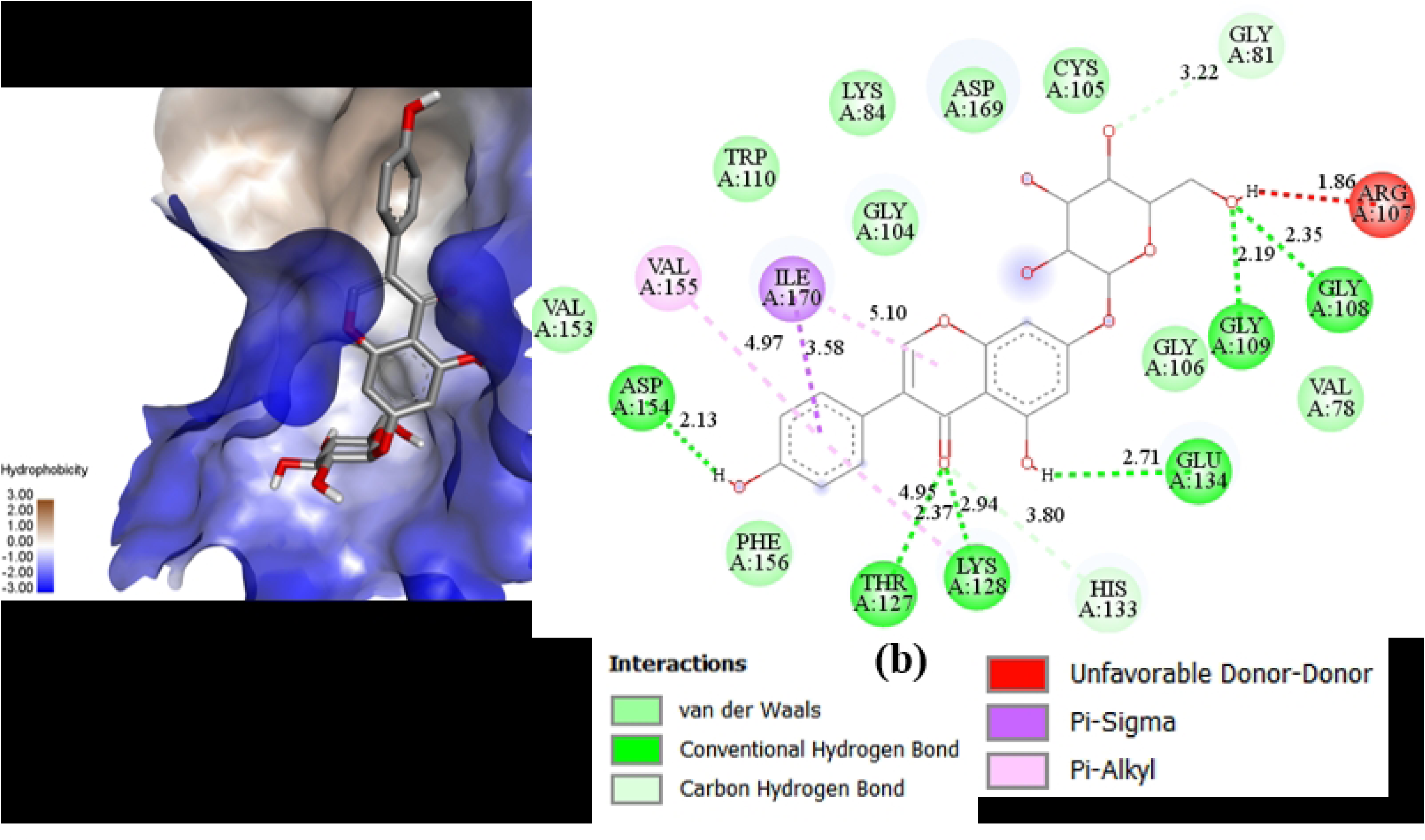
(a) Docking pose of ligand in the cavity with a hydrophobic surface (b) 2D projection of interaction of ligand (FLD1) and protein.

**Fig 5.**
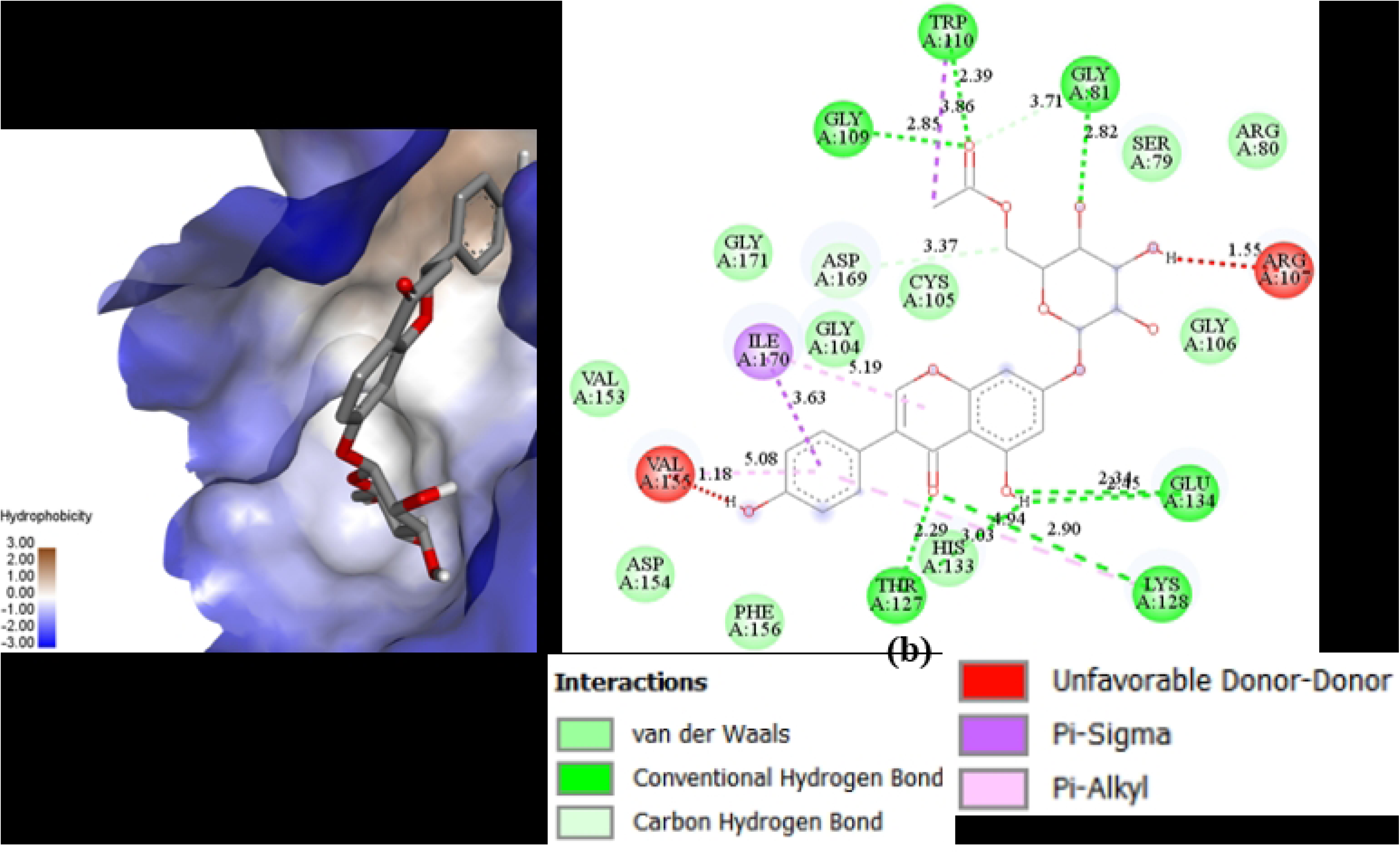
(a) Docking pose of ligand in the cavity with a hydrophobic surface (b) 2D projection of interaction of ligand (FLD2) and protein.

**Fig 6.**
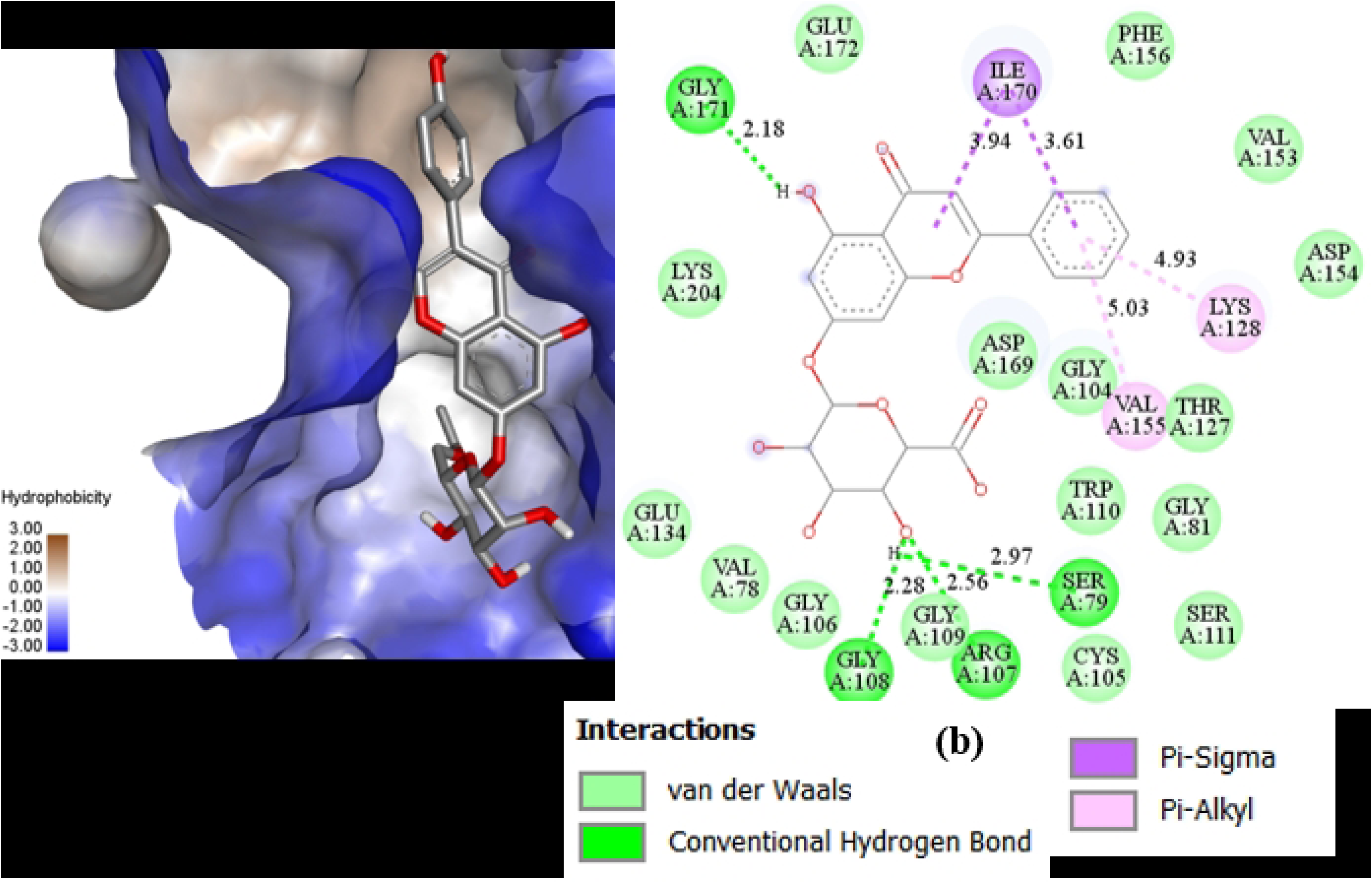
(a) Docking pose of ligand in the cavity with a hydrophobic surface (b) 2D projection of interaction of ligand (FLD3) and protein.

**Fig 7.**
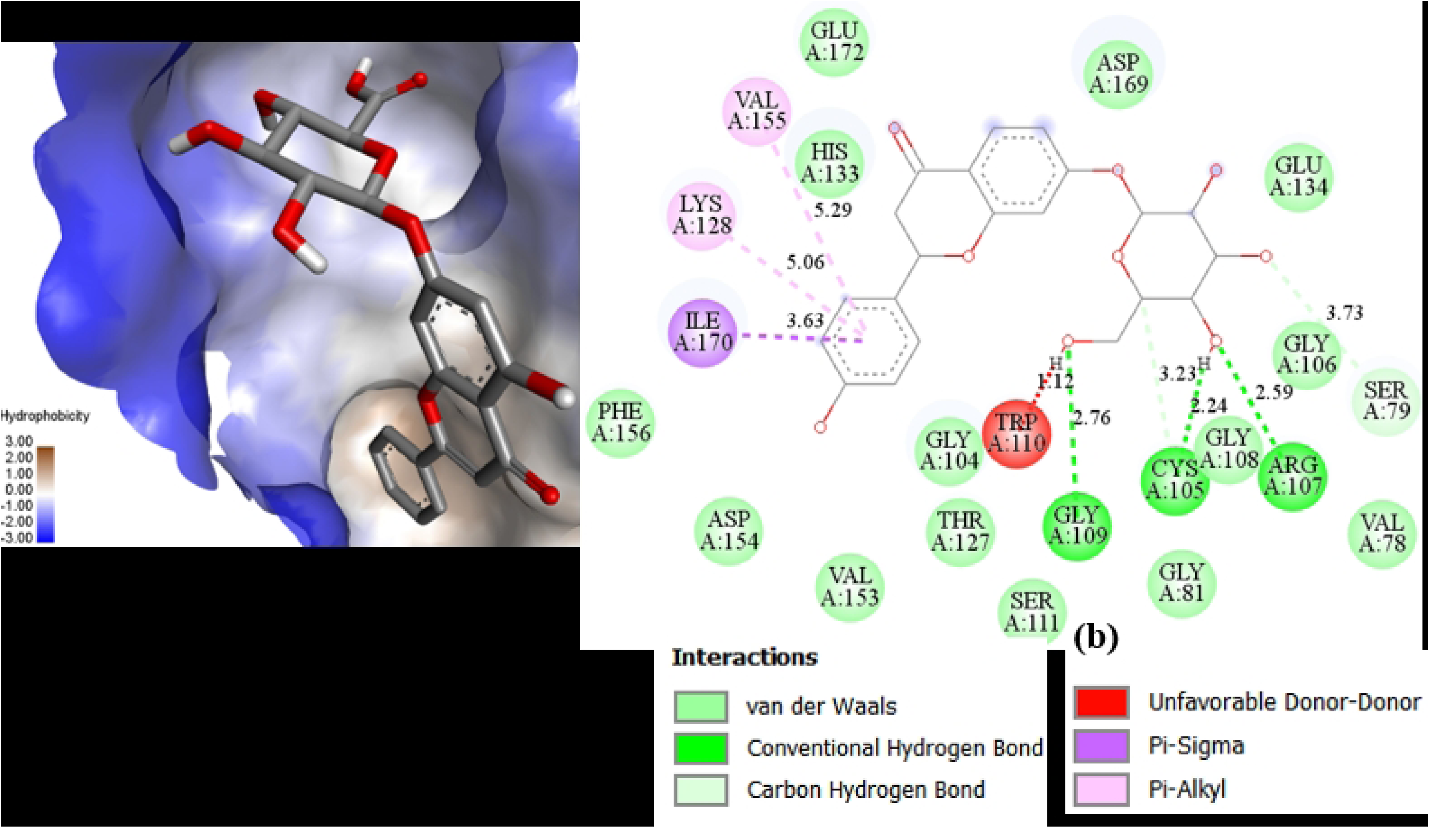
(a) Docking pose of ligand in the cavity with a hydrophobic surface (b) 2D projection of interaction of ligand (FLD4) and protein.

**Fig 8.**
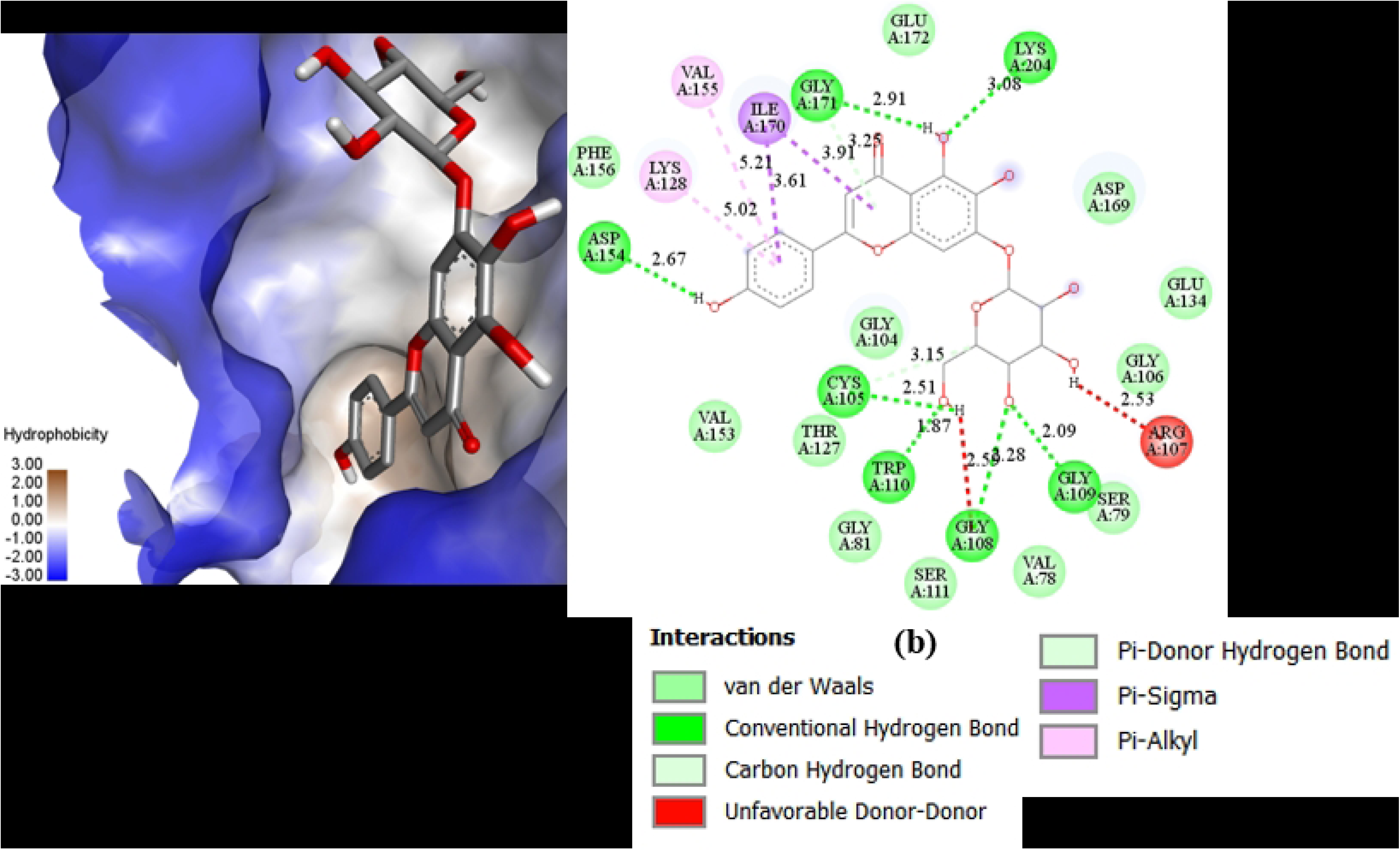
(a) Docking pose of ligand in the cavity with a hydrophobic surface (b) 2D projection of interaction of ligand (FLD5) and protein.

**Table 2:**
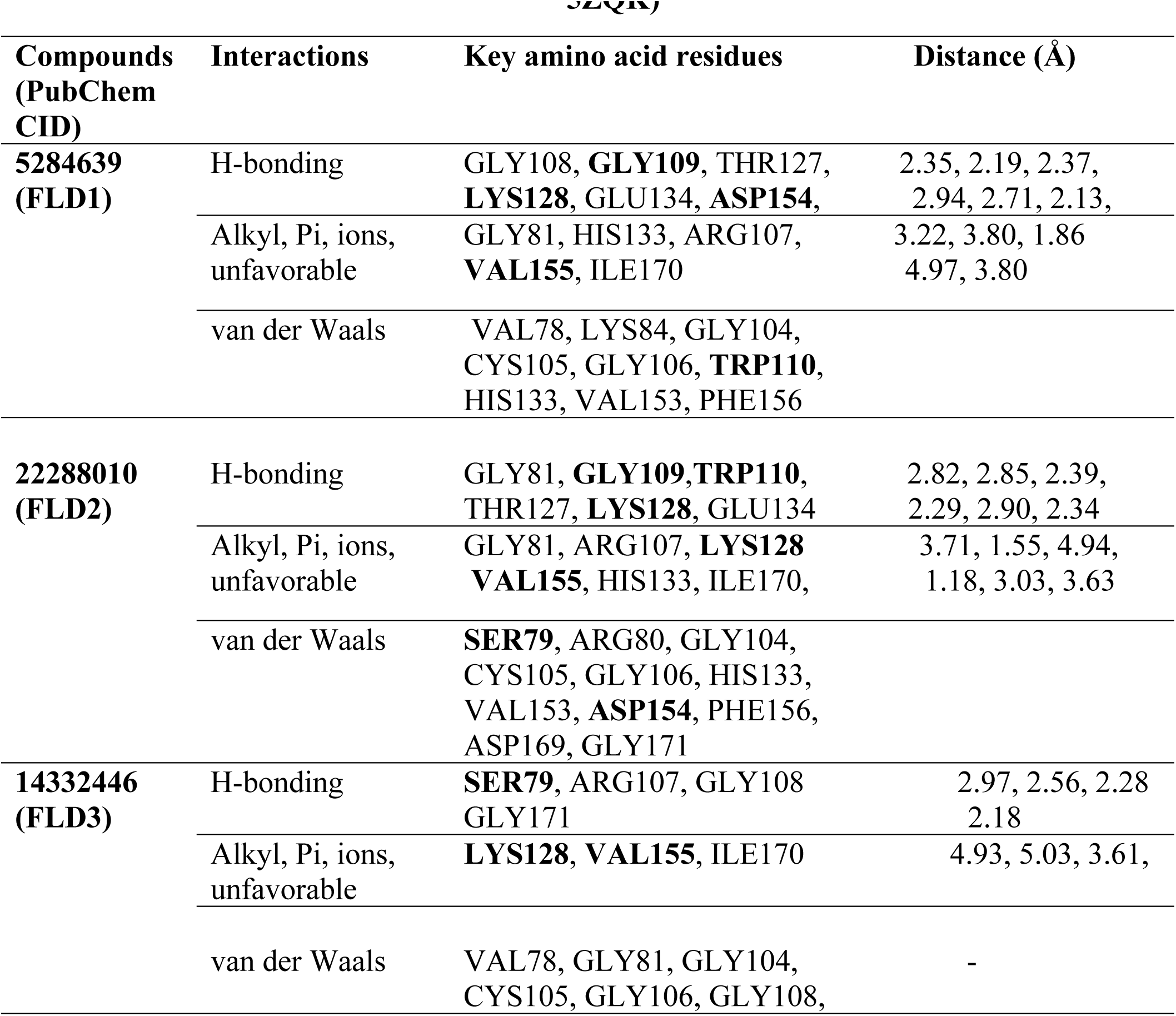
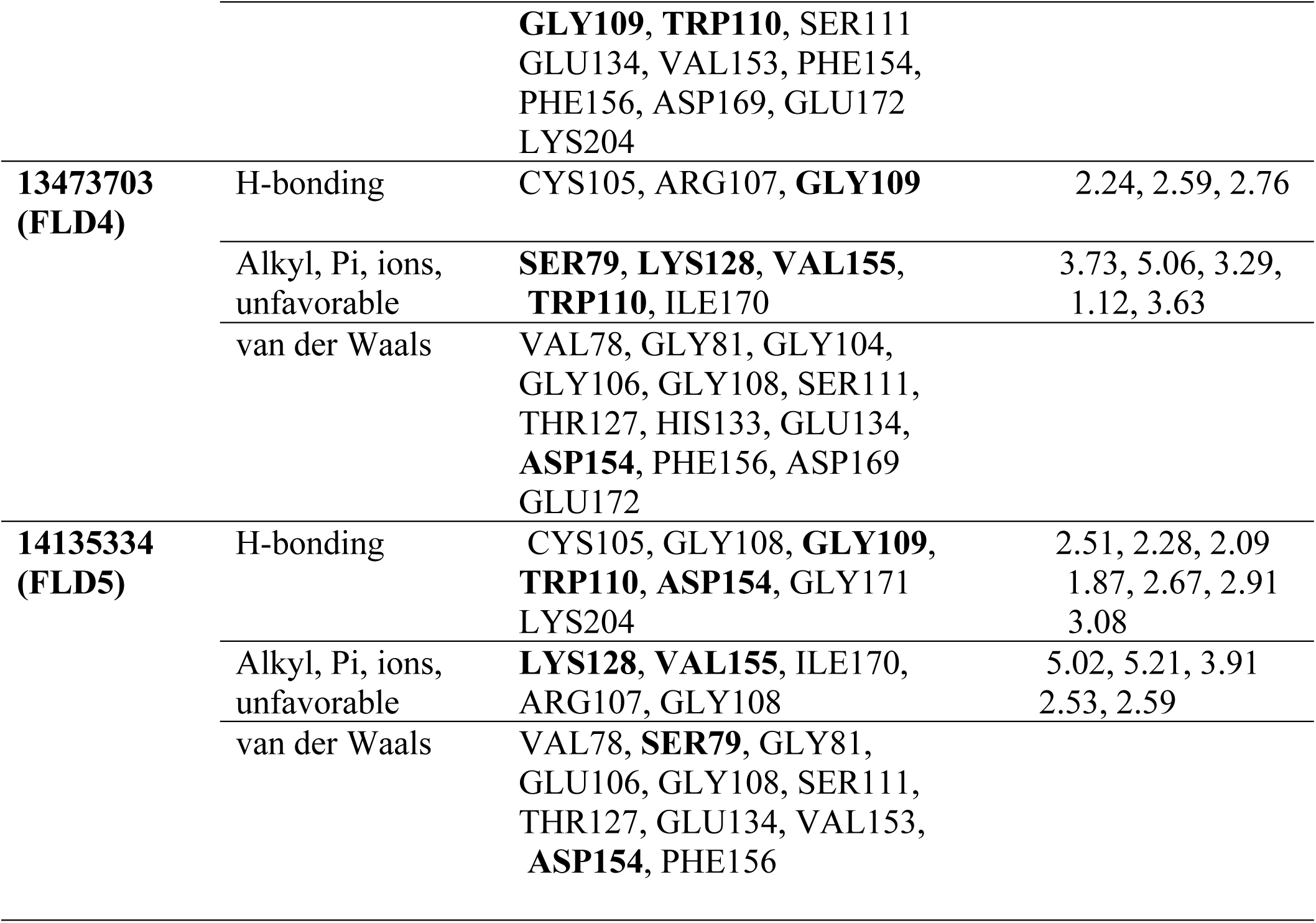
Different types of interaction and key amino acid of ligands with protein (PDB ID 5ZQK)

### 3.6. Molecular dynamics simulation (MDS)

#### 3.6.1 Root mean square deviation (RMSD)

RMSD profiling was used to analyze the dynamic behavior and stability of the protein-ligand complexes [69]. RMSD of the protein relative to the protein backbone and RMSD of the ligand relative to the protein backbone of the different complexes were extracted (Fig 9). The protein backbone exhibited a steady RMSD of *ca.* 3 Å except FLD2 while the RMSD of the ligand relative to the protein backbone varied during the production run. The smooth curve of the protein backbone RMSD indicated that the receptor geometry remained stable upon ligand binding. FLD3, FLD4, and FLD5 showed smooth trajectory at *ca*. 3 Å for 200 ns indicating greater stability of the complexes. RMSD of FLD1 decreased from around 6 Å to 4 Å, with few spikes at 125 ns, and was equilibrated after nearly 130 ns. The RMSD curve at *ca.* 6 Å with several smaller spikes in the RMSD curve of the FLD2 indicated a slightly unstable nature of the complex relative to the other four. A spike at 130 ns of RMSD of protein backbone might be the reason for a bump of RMSD of ligand around the same region of the trajectory. The ligands were stable in the SAM binding pocket during MDS which is also the same as that reported in the previous study [61]. Fig 10 shows snapshots taken at different moments to illustrate how a molecular-level understanding may be attained in terms of the geometry and dynamics of the atoms

**Fig 9.**
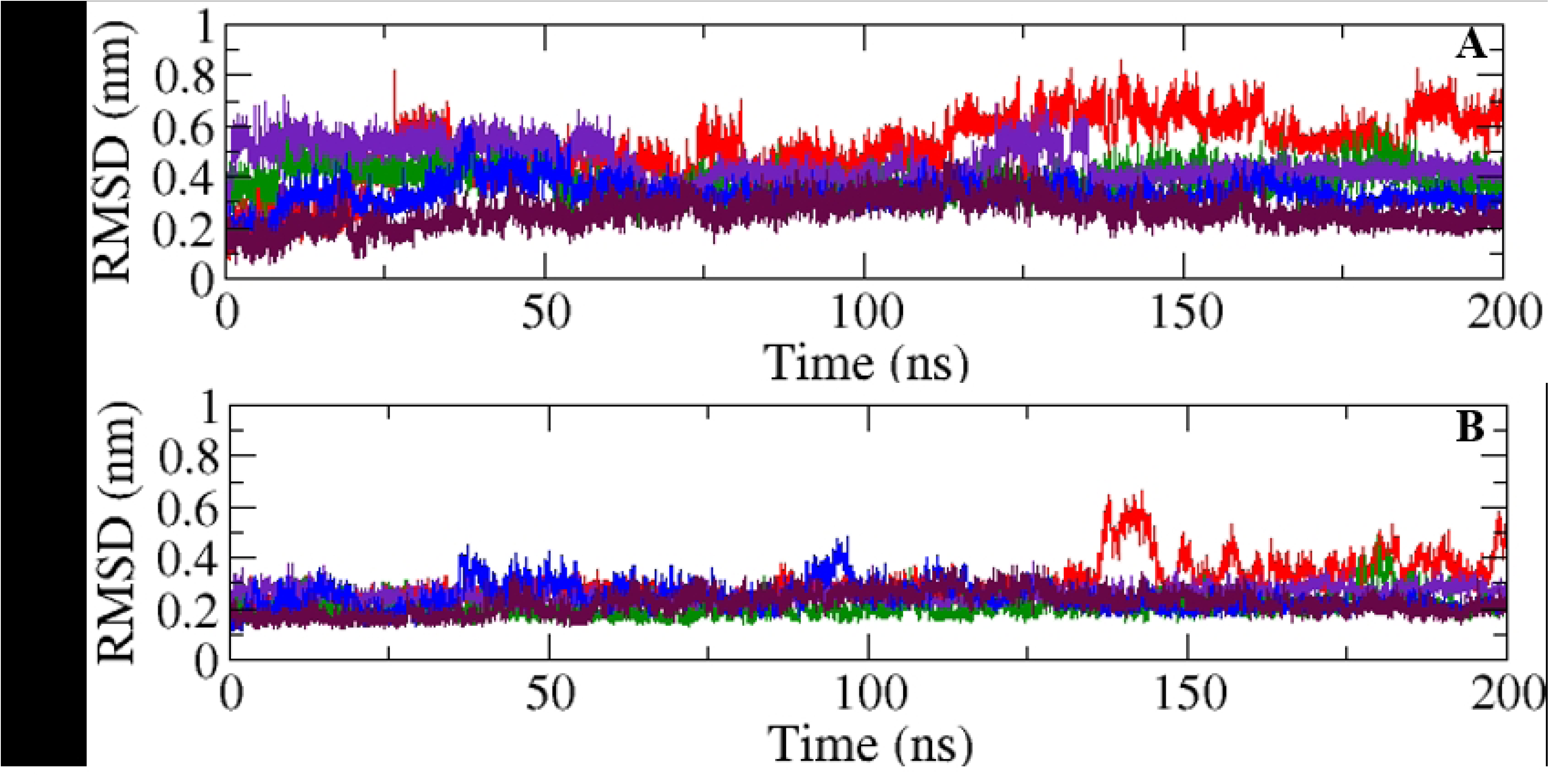
RMSD of (A) ligands and (B) protein backbones both relative to protein backbone in various complexes (protein with flavonoids: Indigo = FLD1, red = FLD2, blue = FLD3, marron = FLD4, green = FLD5)

**Fig 10.**
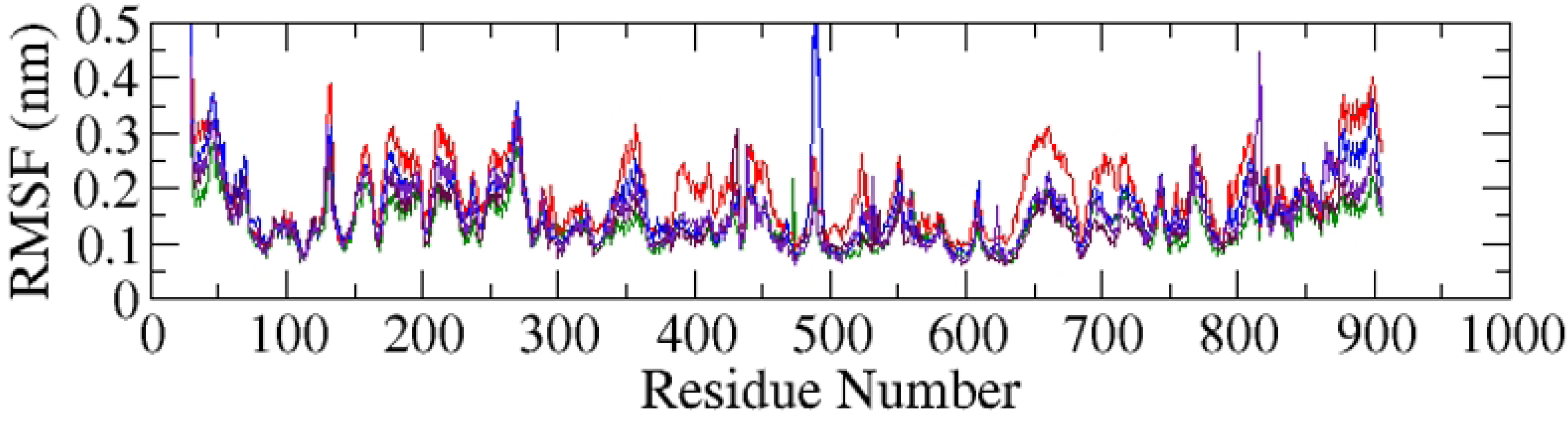
RMSF curves of alpha carbon atoms of protein in protein-ligand complexes for 200 ns MDS trajectory (protein with flavonoids: Indigo curve = FLD1, red curve = FLD2, blue curve = FLD3, marron = FLD4, green4 = FLD5)

#### 3.6.2 Snapshots of ligand at the active site of protein during MDS

Snapshots were retrieved during different instantaneous times in MD simulation to analyze the orientation (rotational motion) and position (translational motion) of the docked ligands. Snapshots at 0, 50, 100, 150, and 200 ns during the simulation, revealed that most of the ligands remained in the same location of active site but with different orientations with few exceptions (Fig 2S). The geometrical behavior at the molecular level can be analyzed to justify the nature of the RMSD curve of ligand relative to the backbone. In the case of FLD1, the ligand shifted farther from the initial position at 50 ns but retrieved its position at the active site with a change in orientation and remained the same along the MD simulations till 200 ns resulting in a lowering of the MD trajectory. In FLD2, the ligand changed its position and orientation significantly at the active site at 100 and 150 ns thus causing few peaks in MD trajectory. The smoothness of the RMSD curve of FLD3 and FLD5 could be justified by the intact position of ligands at all simulation periods. Only minute changes occurred in cases FLD4.

#### 3.6.3 Root mean square fluctuation (RMSF)

The RMSF was calculated from the MDS trajectory for the alpha carbon atom of the protein [70] and was found below 5 Å in most of the complexes indicating the stability of the complexes [71]. In the plot variations in the movement of individual amino acid residues compared to their unbound state. The RMSF of *ca.*4 Å indicated minimal fluctuation in most amino acid residues of the complexes. However, in the protein-FLD3 complex, residues numbered from 475 to 500 exhibited higher fluctuations at *ca*. 5 Å. This is because of the lack of α-helixes or β-sheets and the presence of unbound coils/ loops in the structure [72]. This region does not have significant interaction with the ligand. The minimum fluctuation of the active site residue region at 79 to 155 indicates the stability of the receptor upon ligand binding. It indicated that these fluctuations have a minimal or negligible effect on the destabilization of the complex. The RMSF plot conferred that amino acid residue fluctuations wouldn’t have a major impact on the ligand’s binding in the active pocket. The adduct’s stability would remain unaltered, which would result in an inhibition of regular protein functioning.

#### 3.6.4 Radius of gyration (Rg)

The Rg of the various complexes were extracted from the trajectory and it helps to determine the conformational changes of protein structure. It gives the average distance from the central axis of the macromolecule to all distributed constituents [73]. Rg ranged from 3.17 to 3.3 nm, hinting that the receptor did not undergo significant expansion or contraction during the MD production run except FLD2 (Fig 11). The Rg curve of complex2 shows slight changes after 80 ns. The results imply that there was no considerable receptor expansion or shrinkage in response to ligand binding over the simulated period.

**Fig 11.**
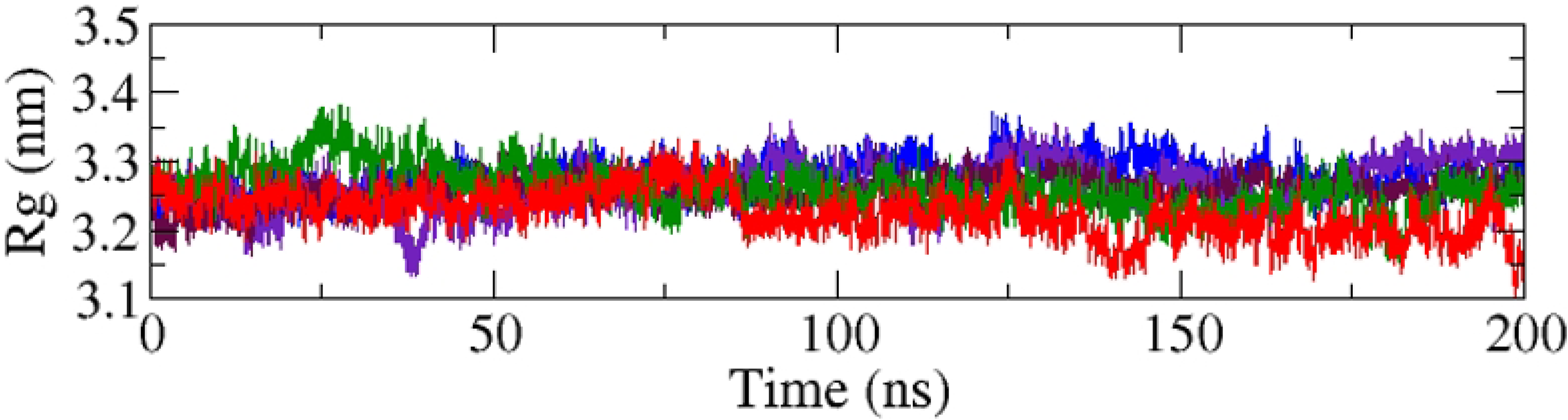
Rg curves of protein in the complex during 200 ns MDS (protein with flavonoids: Indigo curve = FLD1, red curve = FLD2, blue curve = FLD3, marron = FLD4, green = FLD5)

#### 3.6.5 Solvent accessible surface area (SASA)

The SASA helps to determine the total wettable area of the protein. The SASA of the protein was calculated between 390 to 405 nm^2^ (Fig 12) and no notable abrupt changes were observed in the protein structure upon ligand binding. This value implies that during ligand binding, the protein’s hydrophobic portion remains uncovered to the solvent, and its shape remains preserved [74].

**Fig 12.**
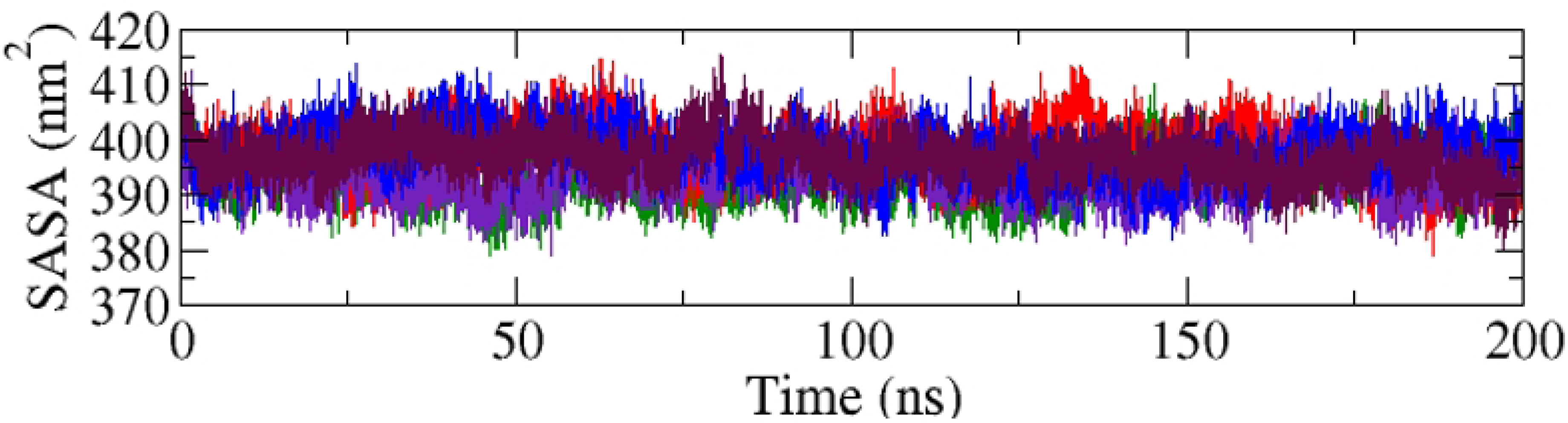
SASA of protein in complexes during 200 ns MDS (protein with flavonoids: Indigo curve = FLD1, red curve = FLD2, blue curve = FLD3, marron = FLD4, green = FLD5)

#### 3.6.6 Hydrogen bond count

The variation of hydrogen bond count during the MDS (Fig 13) plays a pivotal role in the stability of complexes during MDS [75]. The greater the number of hydrogen bonds greater the stability of the complex. The highest hydrogen bond count of up to 8 several times during the MDS was observed in the protein-FLD1 complex (Fig 14). Similarly, the protein-FLD2 complex showed a consistent count of 6 hydrogen bonds. The protein with FLD3 and FLD5 complexes displayed the same hydrogen bond count. In the case of protein with FLD4, a consistent curve in the hydrogen bond count was observed, indicating the relationship between hydrogen bonds and complex stability. The RMSD of protein with FLD1, FLD3, and FLD5 complexes remained below 4 Å and had a minimal translational and rotational motion of ligand, as evidenced by snapshots that’s why the same type of h-bond count was observed. However, the RMSD of the protein with FLD2 is above 4 Å. Slight ligand delocalization was observed in snapshots, and fluctuating hydrogen bond curve during MDS as depicted in snapshots. In complex4, ligands maintain minimal translation and rotational motion, and a corresponding hydrogen bond count was observed. Overall, all parameters indicate the adduct remained stable throughout the simulation.

**Fig 13.**
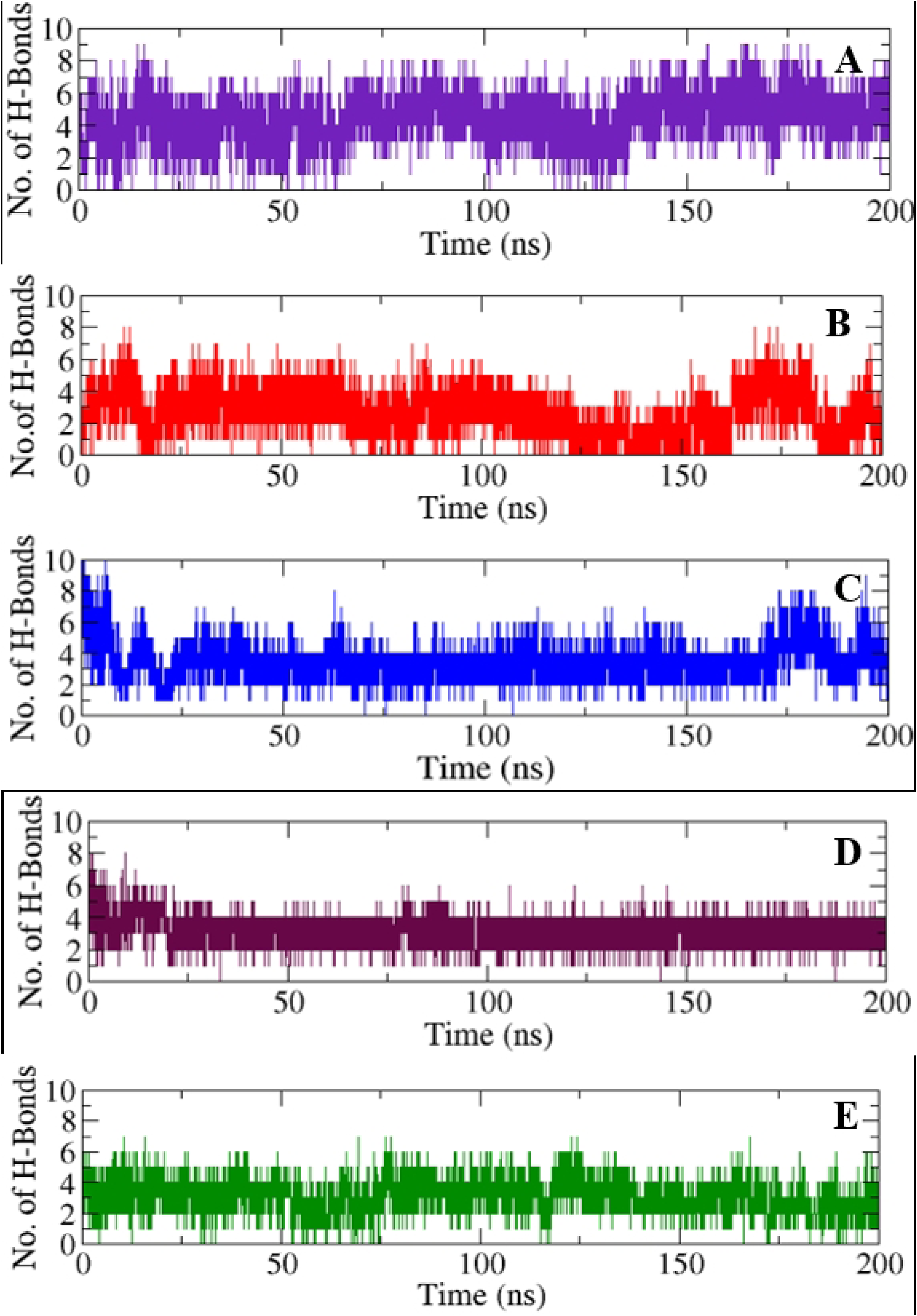
Number of hydrogen bonds between protein and ligand in complexes during 200 ns MDS; A = FLD1 with protein, B = FLD2 with protein, C = FLD3 with protein, D = FLD4 with Protein, E = FLD5 with protein.

### 3.7. Binding free energy changes

The MM/PBSA calculation utilizing the generalized Born model (GBn) was employed to determine the binding free energy difference (ΔG_BFE_) for the equilibrated portion of the trajectory 200 frames (Table 3) [76].

**Table 3.** Change in binding free energies (kcal/mol) of complexes with different components.

| | $\Delta E_{\text{VDWALLS}}$ | $\Delta E_{\text{EL}}$ | $\Delta E_{\text{PB}}$ | $\Delta E_{\text{NPOLAR}}$ | $\Delta G_{\text{GAS}}$ | $\Delta G_{\text{SOLV}}$ | $\Delta G_{\text{BFE}}$ |
| --- | --- | --- | --- | --- | --- | --- | --- |
| FLD1 | -36.07±3.31 | -37.87±12.24 | 43.87±7.54 | -4.3±0.11 | -73.94±10.79 | 39.57±7.54 | -34.38±5.32 |
| FLD2 | -42.37±2.69 | -14.89±16.74 | 44.07±20.72 | -4.56±0.26 | -57.26±16.62 | 39.51±20.52 | -17.75±11.03 |
| FLD3 | -45.67±3.16 | -55.73±15.29 | 67.86±11.88 | -4.47±0.10 | -101.40±15.38 | 63.39±11.84 | -38.01±7.53 |
| FLD4 | -44.95±2.63 | -40.12±5.35 | 56.55±5.62 | -4.39±0.09 | -85.07±5.20 | 52.16±5.60 | -32.91±5.32 |
| FLD5 | -34.49±4.29 | -33.49±14.29 | 50.29±11.10 | -4.77±0.14 | -67.99±12.84 | 45.52±11.06 | -22.47±6.12 |

All the negative values of binding free energy changes (ΔG_BFE_) indicated the feasibility of the adduct formation in all simulated complexes. The highest binding free energy changes were observed with the FLD3-protein complex with −38.01±7.53 kcal/mol, followed by −34.38±5.32, −17.75±11.03, −32.91±5.32 and −22.47±6.12 kcal/mol for the FLD1, FLD2, FLD4, and FLD5 complexes respectively. The frame–by–frame, ΔG_BFE_ of 20 ns of the equilibrated part of the trajectory is depicted (Fig. 3S) and the moving average was calculated after only 50 frames out of 200 frames were calculated. It was always negative at every time for most of the complexes except FLD2. The spontaneous nature of the complex formation indicated the feasibility of inhibition of the receptor. The highest binding free energy dip in the trajectory was around −55 kcal/mol observed in the case of FLD3. Consequently, the interaction between the receptor (5ZQK), and FLD1, FLD2, FLD3, FLD4, and FLD5 were deemed favorable, ensuring stability of the protein-ligand complex throughout the production run.

## 4. Conclusions

The potential of flavonoids to inhibit the NS5 methyl transferase protein of dengue, which is a promising target for dengue virus, has been investigated of possible inhibitory effects of various naturally occurring flavonoids on the NS5 protein of DENV-2 were found from computational methods. Out of the 34 flavonoids examined, 33 exhibited higher binding affinity to the NS5 protein of dengue compared to both reference ligand and an FDA-approved drug. MDS of the top 5 candidates showed good structural stability. The thermodynamical consideration hinted at the spontaneity of the complex formation reaction and maintenance of the ligand pose at the orthosteric site of the receptor. Based on these findings, FLD1, FLD2, FLD3, FLD4, and FLD5 are recommended for additional in vitro and in vivo trials to further warrant their inhibition capabilities against dengue protein.

## Acknowledgments

The authors like to acknowledge Rameshwar Adhikari for his computational lab support.

**Conceptualization:** Achyut Adhikari, Jhashanath Adhikari

**Data curation:** Anuraj Phunyal, Jhashanath Adhikari Subin

**Supervision:** Achyut Adhikari

**Validation:** Anuraj Phunyal

**Writing original draft:** Anuraj phunyal

**Writing-review & editing:** Jhashanath Adhikari Subin, Achyut Adhikari

